# Phylogenomics reveals coincident divergence between giant host sea anemones and the clownfish adaptive radiation

**DOI:** 10.1101/2024.01.24.576469

**Authors:** Aurelien De Jode, Andrea M. Quattrini, Tommaso Chiodo, Marymegan Daly, Catherine S. McFadden, Michael L. Berumen, Christopher P. Meyer, Suzanne Mills, Ricardo Beldade, Aaron Bartholomew, Anna Scott, James D Reimer, Kensuke Yanagi, Takuma Fuji, Estefanía Rodríguez, Benjamin M. Titus

## Abstract

The mutualism between clownfishes (or anemonefishes) and their giant host sea anemones are among the most immediately recognizable animal interactions on the planet and have attracted a great deal of popular and scientific attention [1-5]. However, our evolutionary understanding of this iconic symbiosis comes almost entirely from studies on clownfishes— a charismatic group of 28 described species in the genus *Amphiprion* [2]. Adaptation to venomous sea anemones (Anthozoa: Actiniaria) provided clownfishes with novel habitat space, ultimately triggering the adaptive radiation of the group [2]. Clownfishes diverged from their free-living ancestors 25-30 MYA with their adaptive radiation to sea anemones dating to 13.2 MYA [2, 3]. Far from being mere habitat space, the host sea anemones also receive substantial benefits from hosting clownfishes, making the mutualistic and co-dependent nature of the symbiosis well established [4, 5]. Yet the evolutionary consequences of mutualism with clownfishes have remained a mystery from the host perspective. Here we use bait-capture sequencing to fully resolve the evolutionary relationships among the 10 nominal species of clownfish-hosting sea anemones for the first time (Figure 1). Using time-calibrated divergence dating analyses we calculate divergence times of less than 25 MYA for each host species, with 9 of 10 host species having divergence times within the last 13 MYA (Figure 1). The clownfish-hosting sea anemones thus diversified coincidently with clownfishes, potentially facilitating the clownfish adaptive radiation, and providing the first strong evidence for co-evolutionary patterns in this iconic partnership.

## Main Text

**Figure 1.**
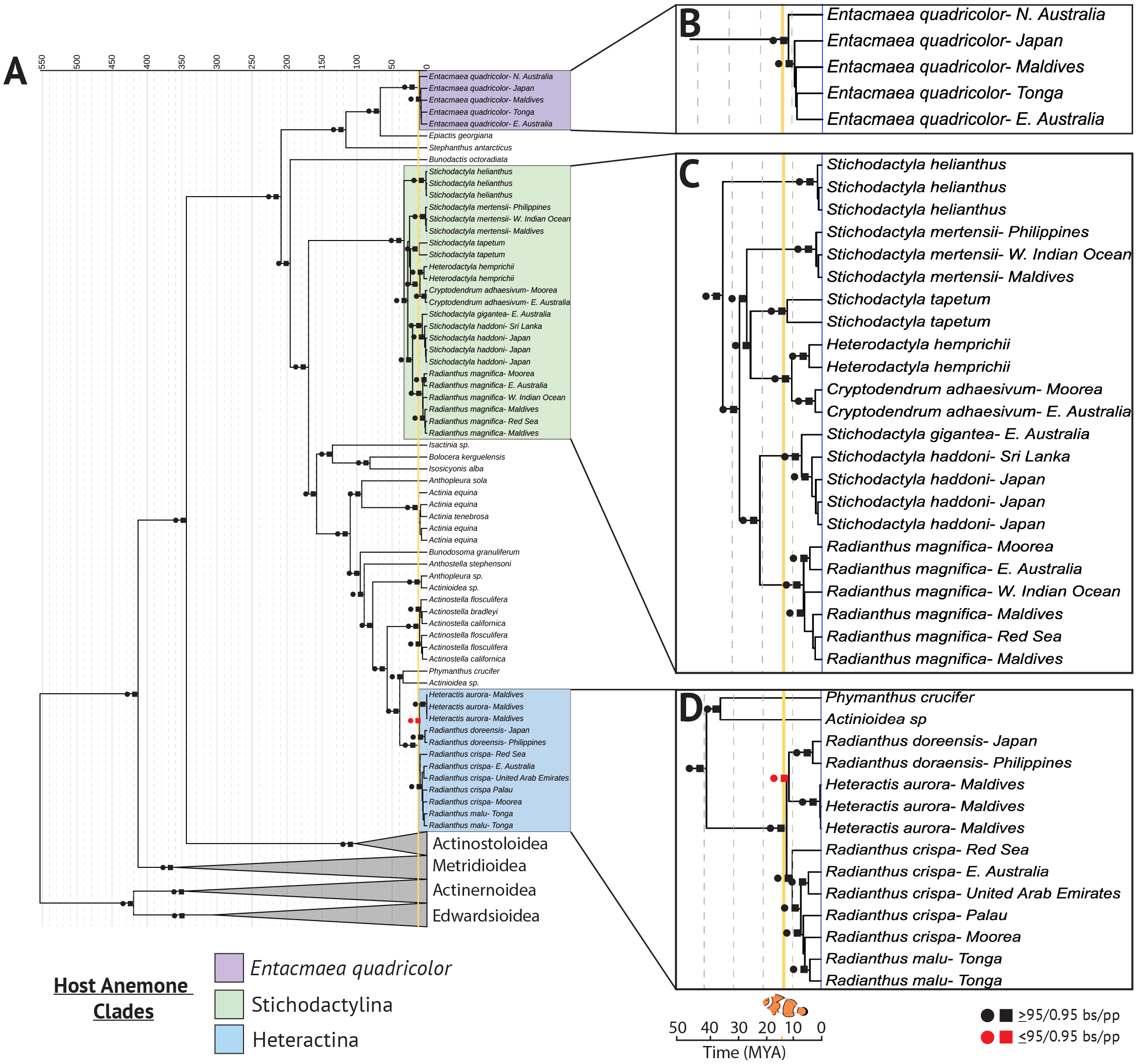
Diversification of the clownfish-hosting sea anemones was coincident with the clownfish adaptive radiation. A) Time-calibrated maximum likelihood cladogram of Order Actiniaria based on 328 ultra-conserved element and exon loci (75% data occupancy matrix). The three clades containing the clownfish hosting sea anemones are highlighted reflecting the multiple evolutionary origins of symbiosis with clownfishes within superfamily Actinioidea. B-D) Detailed time-calibrated maximum likelihood cladograms of *Entacmaea quadricolor*, Clade Stichodactylina, and Clade Heteractina, respectively. In all panels, the orange line denotes the beginning of the clownfish adaptive radiation 13 MYA. Sea anemone superfamilies Actinostoloidea, Metridioidea, Actiniernoidea, and Edwardsioidea are collapsed for clarity.

We assembled a phylogenomic dataset for sea anemones that included representative samples from the 10 clownfish-hosting species (Table S1), which have recently undergone a major nomenclatural revision [5]. Because the clownfish-hosting species have been previously recovered to belong to three clades that are hypothesized to have evolved symbiosis with clownfishes independently [4, 5], we included >50 species of sea anemones across Order Actiniaria to re-evaluate this hypothesis using genomic data (Table S1). Genomic DNA from newly collected samples was extracted, quantified, and prepared for sequencing using bait-capture probes for Class Anthozoa targeting ultra-conserved element (UCE) and exon loci [6]. Final genomic libraries were sequenced using 150bp paired-end sequencing on an Illumina NovaSeq. After sequencing, exon and UCE loci were assembled and extracted from raw sequence data following [6, 7].

Phylogenetic relationships were reconstructed using both maximum likelihood and coalescent-based species tree approaches [8, 9] (see Supplemental Methods). We employed multiple divergence dating analyses to convert phylogenetic trees into ultrametric chronograms and estimate divergence times for the clownfish-hosting sea anemones for the first time [8] (see Supplemental Methods). Minimum and maximum root ages for Actiniaria were set to 424-608 MYA, previously calculated from fossil-calibrated phylogenomic analyses of Anthozoa [7]. Our analyses recovered fully supported phylogenies with nearly identical topologies across Actiniaria (Figure 1A; S1A, S1B). We resolved the 10 described host anemone species to belong to three clades that do not share recent common ancestors—Clades Stichodactylina, Heteractina, and *Entacmaea*—confirming that symbiosis with clownfishes has evolved at least three times independently in sea anemones [4] (Figure 1A). Divergence times for all host anemone species were calculated at less than 25 MYA, with 9 of 10 host species having divergence times between 6.9-11.6 MYA—fully coincident with diversification times calculated for the clownfish adaptive radiation [2, 3] (Figure 1, S2).

Within each clade, we resolve species level relationships for the first time among host taxa. Our analyses calculate the oldest host species, *Stichodactyla mertensii*, to have diverged from its most recent common ancestor 24.9 MYA (95% CI = 21-45.5 MYA; Figure 1, S2). Interestingly, our phylogenetic reconstruction places *S. mertensii* as sister to a clade that contains the non-host anemone *S. tapetum* and members of Family Thalassianthidae. This relationship is fully supported across all analyses making the giant carpet anemones in genus *Stichodactyla*, the largest host anemones, a paraphyletic group (Figure 1C). The remaining carpet anemones, *S. haddoni* and *S. gigantea*, are recovered as sister species that diverged 6.9 MYA (95% CI = 4.2-10.1 MYA) and which form a deeper clade with the magnificent anemone *Radianthus magnifica* dated to 20 MYA (95% CI = 14-27.4 MYA; Figure 1, S2). Although *R. magnifica* is currently considered a single species, our geographic sampling reveals diversity and divergence times within this species that are comparable to the species level divergence seen between *S. haddoni* and *S. gigantea*. We find at least two well resolved sub-clades of *R. magnifica* splitting Indian and Pacific Ocean samples into distinct phylogeographic lineages dated to 7 MYA (95% CI = 4.6-10.1 MYA; Figure 1, S2).

Similarly, the bubble-tip anemone *Entacmaea quadricolor* is recognized as a single widespread species but has been hypothesized to be a cryptic species complex [4]. Our data support this hypothesis, as divergence times between geographic lineages of *E. quadricolor* date to 7.1 MYA (95% CI = 4.2-11.1 MYA; Figure 1, S2). Finally, we date divergence times among the four nominal species within Heteractina (*Heteractis aurora, R. crispa, R. malu, R. doreensis*) to 11.6 MYA (95% CI = 7.8-16.3 MYA; Figure 1, S2). Relationships among species are less well supported within Heteractina, and we recover a sister relationship between *H. aurora* and *R. doreensis* for the first time (Figure 1, S1A, S1B). Like *E. quadricolor* and *R. magnifica*, we find undescribed diversity within *R. crispa*, with lineages from the Arabian Peninsula diverging from the rest of the Indo-West Pacific 9.8 MYA (95% CI = 6.7-14.1 MYA; Figure 1, S2), followed by geographic lineages from Japan, Palau, Australia, and Moorea all variously dated between 5-7 MYA. Interestingly, we find *R. malu* as a distinct lineage, but nested within the broader *R. crispa* complex, casting doubt on the taxonomic status of this putative species (Figure 1, S1, S2).

Mutualisms are tightly linked ecological interactions that are expected to impact the formation and distribution of species level biodiversity among constituent partners [10]. Our data are consistent with these expectations. The ecological setting that led to the evolution of the clownfish-sea anemone symbiosis has been a long-standing mystery. Following divergence from their free-living relatives, over 15 million years passed before clownfishes adaptively radiated to sea anemones throughout the Indo-West Pacific. Why? Our findings suggest that most of the host anemone diversity hadn’t evolved yet. We hypothesize that the coincident diversification between host anemones and clownfishes beginning ~13 MYA ultimately facilitated the clownfish adaptive radiation. Our results provide the first strong evidence for co-evolutionary patterns between clownfishes and their host sea anemones, and point to the importance of understanding host sea anemone biology for a comprehensive understanding of this model mutualism.

## Supporting information

Supplemental information contains one table, two figures, detailed methods, and author contributions.

## Acknowledgements

We thank the Small Island Research Station (Fares-Maathoda, Maldives) and CRIOBE (Moorea, French Polynesia) for field research support and logistics, especially Mohamed Aslam, Ali Zahir, and Rahula Suhail. Kevin Kohen (Live Aquaria) and Laura Simmons (Cairns Marine) provided anemone samples from Tonga and Australia. Lily Berniker (AMNH) helped with sample handling and accession.

This research was funded by National Science Foundation (NSF) award DEB-1934274 to BMT and ER, NSF award DEB-145781 to ER, and DEB-1457817 to CSM, The University of Alabama Start-up funds to BMT, the Gerstner Scholars Postdoctoral Fellowship and Gerstner Family Foundation, the Lerner Gray Fund for Marine Research, and Richard Gilder Graduate School at the American Museum of Natural History to BMT. Sample collection from Panama was supported by NSF award DEB-1456196 to ER. Fieldwork in Palau by JDR was part of the SATREPS P-CoRIE Project “Sustainable management of coral reef and island ecosystem: responding to the threat of climate change” funded by the Japan Science and Technology Agency (JST) and the Japan International Cooperation Agency (JICA) in cooperation with Palau International Coral Reef Center and Palau Community College. Fieldwork in Japan was funded by the Japan Society for the Promotion of Science (JSPS) Kakenhi Grants (JP255440221 to KY, and JP17K15198 and JP17H01913 grants to TF), and Kagoshima University adopted by the Ministry of Education, Culture, Sports, Science and Technology, Japan (Establishment of Research and Education Network of Biodiversity and its Conservation in the Satsunan Islands project to TF). Fieldwork in Saudi Arabia was provided in part by the King Abdullah University of Science and Technology (KAUST) Office of Competitive Research Funds under Award No. CRG-1-2012-BER-002 and baseline research funds to MLB.

## Inclusion and Diversity

We support inclusive, diverse, and equitable conduct of research

