## Supplemental information contains one table, two figures, detailed methods, and author contributions. for "Phylogenomics reveals coincident divergence between giant host sea anemones and the clownfish adaptive radiation"

**Table S1.** Sea anemone samples sequenced and included in this study. Table includes information on Superfamily, Genus, and Species designations based on the currently accepted taxonomy. Sample size per species and per locality is denoted by *N.* Locality indicates country of origin for each sample. Dataset refers to the bait-capture dataset that sample is included in: 1) Order Actiniaria, 2) Clade Stichodactylina, 3) Clade Heteractina. Accession Numbers (Accession #) in bold are new to this study.

| Superfamily | Genus | Species | *N* | Locality | Accession # |
| --- | --- | --- | --- | --- | --- |
| Actinernoidea | *Synhalcurias* | *elegans* | 1 | Japanese Archipelago | SAMN13244953 |
| Actinioidea | *Actinostella* | *californica* | 2 | Baja, Mexico | TBD-TBD |
| Actinioidea | *Actinostella* | *bradleyi* | 1 | Baja, Mexico | TBD |
| Actinioidea |  | *flosculifera* | 1 | Bocas del Toro, Panama | TBD |
| Actinioidea |  | *flosculifera* | 1 | Unknown | TBD |
| Actinioidea |  | sp. | 1 | Bocas del Toro, Panama | TBD |
| Actinioidea | *Actinioidea* | sp. | 1 | Panama | SAMN07774921 |
| Actinioidea | *Actinia* | *equina* | 2 | Spain | SAMN13244941 |
| Actinioidea |  | *equina* | 1 | Iceland | TBD |
| Actinioidea |  | *equina* | 1 | Ireland | TBD |
| Actinioidea | *Actinia* | *tenebrosa* | 1 | New Zealand | TBD |
| Actinioidea | *Anthopleura* | sp. | 1 | Panama | SAMN13244886 |
| Actinioidea | *Anthostella* | *stephensoni* | 1 | South Africa | SAMN13244945 |
| Actinioidea | *Bolocera* | *kerguelensis* | 1 | South Orkneys, Antarctica | SAMN13244888 |
| Actinioidea | *Bunodactis* | *octoradiata* | 1 | Patagonia, Chile | TBD |
| Actinioidea | *Bunodosoma* | *granuliferum* | 1 | Panama | SAMN13244947 |
| Actinioidea | *Entacmaea* | *quadricolor* | 1 | Southern Great Barrier Reef, Australia | TBD |
| Actinioidea |  | *quadricolor* | 1 | Northern Territories,  Australia | TBD |
| Actinioidea |  | *quadricolor* | 1 | Fares-Maathoda, Maldives | TBD |
| Actinioidea |  | *quadricolor* | 1 | Japanese Archipelago | TBD |
| Actinioidea |  | *quadricolor* | 1 | Tonga | TBD |
| Actinioidea | *Epiactis* | *georgiana* | 1 | South Orkneys, Antarctica | SAMN13244887 |
| Actinioidea | *Isactinia* | sp. | 1 | New Zealand | TBD |
| Actinioidea | *Isosicyonis* | *alba* | 1 | Antarctica | SAMN07774928 |
| Actinioidea | *Stephanthus* | *antarcticus* | 1 | Antarctica | SAMN13244952 |
| Actinioidea | *Heteractis* | *aurora* | 3 | Fares-Maathoda, Maldives | TBD |
|  |  | *aurora* | 1 | Japanese Archipelago | TBD |
| Actinioidea | *Phymanthus* | *crucifer* | 1 | Panama | SAMN13244890 |
| Actinioidea | *Radianthus* | *crispa* | 3 | Thuwal, Saudi Arabia | TBD |
| Actinioidea |  | *crispa* | 3 | Gulf of Oman,  United Arab Emirates | TBD |
| Actinioidea |  | *crispa* | 4 | Moorea, French Polynesia | TBD |
| Actinioidea |  | *crispa* | 3 | Southern Great Barrier Reef,  Australia | TBD |
| Actinioidea |  | *crispa* | 2 | Tonga | TBD |
| Actinioidea |  | *crispa* | 3 | Palau | TBD |
| Actinioidea |  | *crispa* | 7 | Japanese Archipelago | TBD |
| Actinioidea | *Radianthus* | *doreensis* | 2 | Philippines | TBD |
| Actinioidea |  | *doreensis* | 8 | Japanese Archipelago | TBD |
| Actinioidea | *Radianthus* | *magnifica* | 1 | Moorea, French Polynesia | TBD |
| Actinioidea |  | *magnifica* | 1 | Northern Great Barrier Reef, Australia | TBD |
| Actinioidea |  | *magnifica* | 1 | Scattered Islands | TBD |
| Actinioidea |  | *magnifica* | 1 | Thuwal, Saudi Arabia | TBD |
| Actinioidea |  | *magnifica* | 2 | Fares-Maathoda, Maldives | TBD |
| Actinioidea | *Radianthus* | *malu* | 2 | Tonga | TBD |
| Actinioidea | *Stichodactyla* | *gigantea* | 1 | Northern Great Barrier Reef,  Australia | TBD |
| Actinioidea | *Stichodactyla* | *haddoni* | 2 | Sri Lanka | TBD |
| Actinioidea |  | *haddoni* | 8 | Japanese Archipelago | TBD |
| Actinioidea | *Stichodactyla* | *helianthus* | 5 | Bocas del Toro, Panama | SAMN13244891-TBD |
| Actinioidea | *Stichodactyla* | *mertensii* | 2 | Fares-Maathoda, Maldives | TBD |
|  |  | *mertensii* | 1 | Phillipines |  |
| Actinioidea |  | *mertensii* | 1 | Scattered Islands | TBD |
| Actinioidea |  | *mertensii* | 1 | Thuwal, Saudi Arabia | TBD |
| Actinioidea |  | *mertensii* | 2 | Japanese Archipelago | TBD |
| Actinioidea | *Stichodactyla* | *tapetum* | 1 | Northern Great Barrier Reef,  Australia | TBD |
| Actinioidea |  | *tapetum* | 1 | Vietnam | TBD |
| Actinioidea | *Cryptodendrum* | *adhaesivum* | 1 | Northern Great Barrier Reef,  Australia | TBD |
| Actinioidea |  | *adhaesivum* | 1 | Moorea, French Polynesia | TBD |
| Actinioidea |  | *adhaesivum* | 1 | Fares-Maathoda, Maldives | TBD |
| Actinioidea | *Thalassianthus* | *hemprichii* | 2 | Northern Great Barrier Reef,  Australia | TBD |
| Actinioidea | *Unknown* | sp*.* | 1 | Unknown | TBD |
| Actinostoloidea | *Actinostola* | sp. | 1 | South Orkneys, Antarctica | SAMN13244889 |
| Actinostoloidea | *Antholoba* | *achates* | 1 | Patagonia, Chile | SAMN13244944 |
| Actinostoloidea | *Stomphia* | *didemon* | 1 | Washington, USA | SAMN13244908 |
| Actinostoloidea | *Halcampella* | *fasciata* | 1 | South Orkneys, Antarctica | SAMN13244892 |
| Actinostoloidea | *Sicyonis* | sp. | 1 | Antarctica | SAMN07774926 |
| Edwardioidea | *Edwardsia* | *timida* | 1 | Ireland | SAMN13244884 |
| Edwardioidea | *Nematostella* | *vectensis* | 1 | Unknown | SAMN02953687 |
| Metridioidea | *Exaiptasia* | *diaphana* | 1 | Unknown | SAMN03839803 |
| Metridioidea | *Laviactis* | *lucida* | 1 | Panama | SAMN13244950 |
| Metridioidea | *Alicia* | *sansibarensis* | 1 | South Africa | SAMN13244943 |
| Metridioidea | *Lebrunia* | *neglecta* | 1 | Bocas del Toro, Panama | TBD |
| Metridioidea | *Antipodactis* | *awii* | 1 | Antarctica | SAMN13244946 |
| Metridioidea | *Bunodeopsis* | *globulifera* | 1 | Bocas del Toro, Panama | TBD |
| Metridioidea | *Diadumene* | *leucolena* | 1 | Brazil | SAMN13244966 |
| Metridioidea | *Galatheanthemum* | *cf. profundale* | 1 | Japanese Archipelago | SAMN13244949 |
| Metridioidea | *Halcurias* | *pilatus* | 1 | Chile | SAMN07774925 |
| Metridioidea | *Actinauge* | *richardi* | 1 | Ireland | SAMN13244965 |
| Metridioidea | *Phelliactis* | sp. | 1 | Ireland | SAMN13244964 |
| Metridioidea | *Metridium* | *senile* | 1 | Maine, USA | SAMN13244909 |
| Metridioidea | *Sagartia* | *troglodytes* | 1 | Ireland | SAMN13244951 |

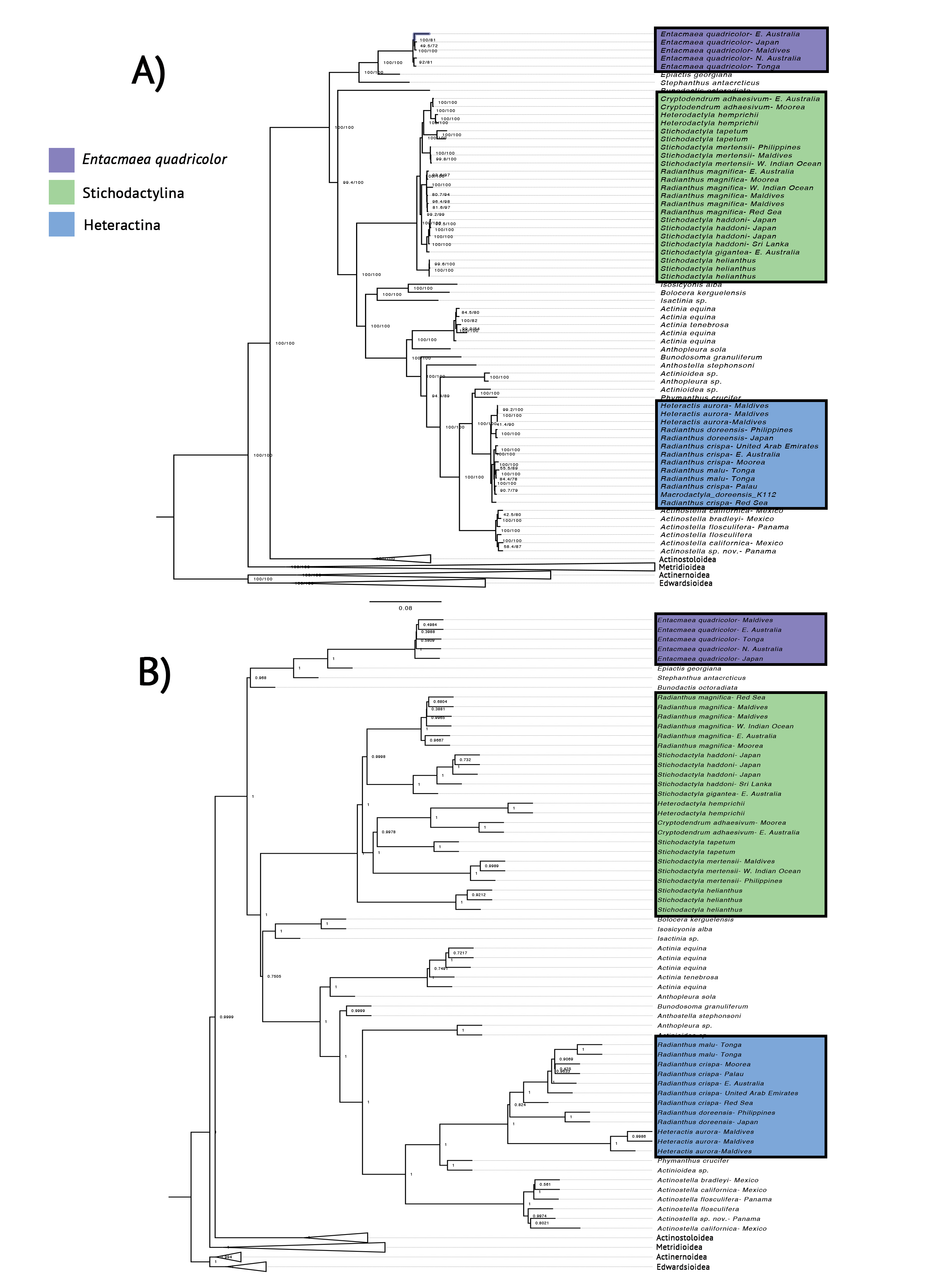

Figure S1. A) Maximum likelihood (IQTREE-2) and B) ASTRAL species tree reconstruction of the clownfish-hosting sea anemones (purple, green, and blue highlighted clades). Trees are based on 75% occupancy matrix of 328 ultra-conserved element and exon loci. Branch lengths for ML tree are presented in substitutions per site. Node support values for ML tree presented from ultrafast bootstrap support and sh-like approximate likelihood ratio tests. Node support values for ASTRAL species tree presented as posterior probabilities.

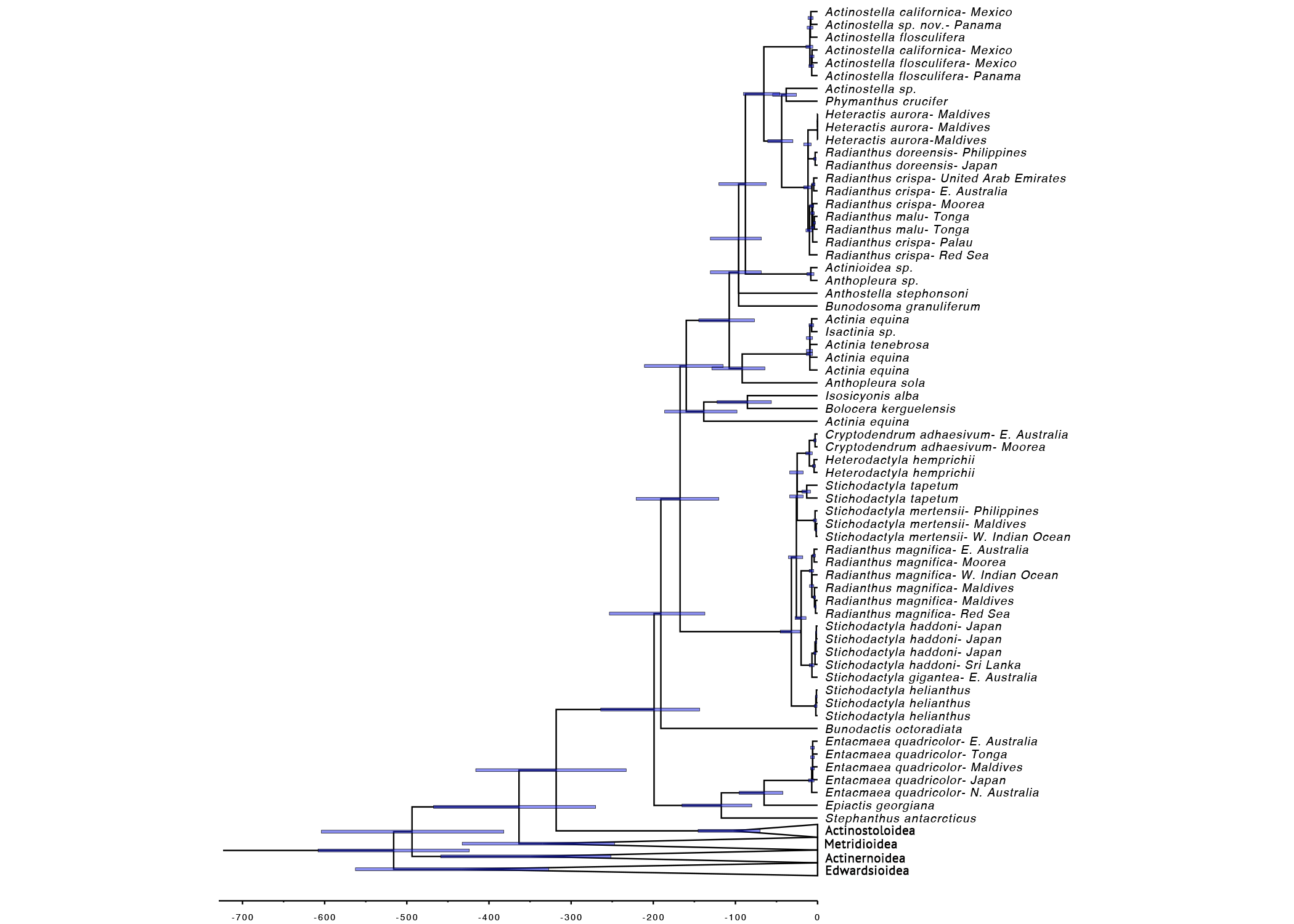

Figure S2. Time-calibrated maximum likelihood cladogram of Order Actiniaria based on 328 ultra-conserved element and exon loci (75% data occupancy matrix). Blue bars on each node represent 95% confidence intervals from divergence dating analyses in IQTREE-2.

**Supplemental Methods**

***Taxonomic sampling:*** Representative samples from all 10 species of clownfish-hosting sea anemones were newly collected and sequenced in this study (Table S1). Samples were acquired through a combination of field collected tissue samples or sourced via the ornamental aquarium trade directly from specific sample localities. Field samples were photographed prior to sampling and collected underwater by hand using forceps and scissors on SCUBA. Tissue samples were placed into separately labeled whirl-pack bags and preserved using 95% EtOH or RNAlater. Where possible, we obtained samples for each species from multiple sample localities throughout the Indo-West Pacific. In total, N = 33 clownfish-hosting sea anemone samples were included and sequenced in this study. We also included samples from non-host sea anemones within clade Stichodactylina, Superfamily Actinioidea, and across Order Actiniaria broadly to represent as much sea anemone diversity as possible (Table S1). These samples and data were obtained through a combination of field collection, museum holdings, ornamental aquarium trade, and publicly available UCE data previously published by Quattrini et al. (2018, 2020; Table S1). Finally, we included UCE data from the sea anemones *Anthopleura sola, Exaiptasia diaphana,* and *Nematostella vectensis* by mining UCE loci out of publicly available genomes (Putnam et al. 2005; Baumgarten et al. 2015; Cornwell et al. 2022; Fletcher et al. 2022) following the bioinformatic pipeline in phyluce v1.7.1 (Faircloth 2016). In total, our final dataset included N = 86 sea anemone samples.

***DNA extraction, library preparation, and sequencing*:** For each newly collected sample, total genomic DNA was extracted using Qiagen DNeasy blood and tissue spin-column kits (Qiagen Inc.) and stored at -20ºC. Prior to sequencing, DNA quantity (ng/µL) and quality (260/280) were assessed using a Qubit 2.0 fluorometer and NanoDrop spectrophotometer, respectively. Samples were prepared and sequenced across three individual Illumina libraries. Sequence-capture library preparation was performed at both Harvey Mudd College and Arbor BioSciences (Ann Arbor, MI) following the protocol developed by Quattrini et al. (2018). We used the Hexacorallia specific bait set from Cowman et al. (2020), which was further modified to specifically target actiniarians (see Glon et al. 2021). The resulting bait set contained 17,268 baits targeting 2496 loci (e.g. exons and UCEs), which were synthesized by Arbor BioSciences.

For each sample, up to 1000ng of genomic DNA was carried forward for library preparation, which was sheared to a target fragment size of 400-800bp using a Covaris Ultrasonicator. Library preparation was performed using a Kapa Hyper Prep Kit for bait-capture sequencing, with universal Y-yoke oligonucleotide adapters and iTru dual-indexed primers (Glenn et al. 2016). Libraries were pooled into equimolar ratios (100ng) and target enrichment was performed using MyBaits v.IV protocol using a 500 ng/rxn concentration of baits (Glon et al. 2021). Bait-capture enriched libraries were sequenced at Arbor BioSciences on an Illumina NovaSeq using 150bp paired-end sequencing.

***UCE dataset assembly:*** After sequencing, exon and UCE loci were extracted and assembled from raw sequence data following Quattrini et al. (2018, 2020) using the hexa_v2_final bait set (see Glon et al. 2021). For each sample, raw sequence reads were first cleaned using Illumiprocessor v2.1.0 in phyluce (Faircloth 2016) and assembled using SPAdes v3.14.1 (Bankevich et al. 2012) with the -careful and -cov-cutoff 2 parameters. We then used phyluce, as described in online tutorials to extract exon and UCE loci and assemble the final dataset. For our final dataset we further created 75% and 50% taxon-occupancy matrices for each locus to explore how missing data threshold impacted the number of recovered loci and phylogenetic reconstruction at various hierarchical levels. Each data matrix was aligned using MAFFT v7.475 (Katoh et al. 2002), internally trimmed using gblocks v0.91b with default parameters and concatenated using phyluce into a single large super matrix.

***Phylogenomic reconstruction and divergence dating analyses:***

Phylogenomic analyses were conducted using Maximum Likelihood reconstruction as implemented in IQ-TREE 2 (Minh et al. 2020). We ran partitioned analyses for each concatenated dataset by using ModelFinder (Kalyaanamorrthy et al. 2017) to find the best fit nucleotide substitution model per locus. In IQTree we conducted ultrafast bootstrapping (Minh et al., 2013) and Sh-like approximate likelihood ratio tests (Guindon et al. 2010) to evaluate the strength of nodal support for each analysis. In order to capture as many UCE and exon loci as possible, we opted to use *N. vectensis* as our outgroup taxa for our full dataset of Order Actiniaria, rather than a non-actiniarian, as our focus here is focused on relationships within superfamily Actinioidea rather than across the order. Resulting trees were visualized in FigTree 1.4.4 (Rambaut 2014)

Next, we conducted species tree phylogenetic analyses using ASTRAL III (Zhang et al. 2017) and CASTER (Zhang et al. 2023) to provide an alternative to our concatenated maximum likelihood reconstruction in IQ-TREE 2. Both ASTRAL III and CASTER are coalescent-based species tree methods that estimates species trees from multiple genes under the multi-species coalescent model. Thus, both approaches account for incomplete lineage sorting, a biological process that is especially common at shallow evolutionary timescales. ASTRAL and CASTER differ slightly in their approach. ASTRAL estimates species trees given a set of input gene trees (Zhang et al. 2017). Gene trees for each locus are first constructed in IQ-TREE2 using the best fit model of nucleotide evolution selected in ModelFinder. ASTRAL analyses were conducted with astral-hybrid v1.15.2.3. Species trees were visualized in FigTree. In contrast, CASTER estimates species trees directly from nucleotide alignments. For each locus, we estimated species trees in CASTER v1.15.01 using UCE alignments that were converted into fasta file format using phyluce. Species trees in CASTER were estimated using both HKY and GTR models of nucleotide substitution. In both ASTRAL and CASTER we conducted analyses for each data matrix (e.g. 50% and 75% occupancy).

To estimate divergence times among the clownfish hosting anemones, we used Chronos and IQ-TREE 2. Both approaches fit a chronogram to a phylogenetic tree whose branch lengths are in substitutions per site but do so in different ways. We used our 75% occupancy phylogenetic tree as our input guide tree for both analyses. Chronos conducts molecular dating by penalized likelihood and maximum likelihood given a known minimum and maximum root age. We used Chronos in the ape package in R and set the root minimum and maximum to 424-608 MYA, previously calculated from fossil-calibrated phylogenomic analyses of Anthozoa (Quattrini et al. 2020). We used a “clock” model, set the lambda parameter to 1, and kept default chronos.control settings. In IQ-TREE 2, molecular dating analyses employs a least squares dating (LSD2) method. Like Chronos we provided the same guide tree and minimum and maximum root ages for Actinaria. In IQ-TREE we also calculated 95% confidence intervals for divergence time estimates.
